# gVCF2CNV: a scalable pipeline for CNV detection from whole-genome sequencing data

**DOI:** 10.64898/2026.09.02.748565

**Authors:** Mame Seynabou Diop, Florian Bénitière, Kuldeep Kumar, Benjamin Clark, Martineau Jean-Louis, Zohra Saci, Guillaume Huguet, Sylvie Hamel, Sébastien Jacquemont

**Affiliations:** Université de Montréal, 2900 Bd Édouard-Montpetit, Montréal, QC, H3T 1J4; Centre de recherche CHU Sainte-Justine, 3175 Chemin de la Côte-Sainte-Catherine, Montréal, QC, H3T 1C5, Canada; Département d’Informatique et de Recherche Opérationnelle, Université de Montréal, 3150 Rue Jean-Brillant, Montréal, QC, H3T 1N8, Canada

## Abstract

**Motivation:** Copy-number variants (CNVs) contribute to human disease and population trait variation. CNV detection from large whole-genome sequencing cohorts remains computationally demanding, as most methods require BAM or CRAM files. Genomic VCF (gVCF) files are smaller, routinely generated by standard variant-calling workflows, and contain the read depth and allelic information needed for CNV detection. However, gVCF files are not directly compatible with established CNV callers that rely on Log R Ratio (LRR) and B Allele Frequency (BAF) signals.

**Results:** We present gVCF2CNV, a Nextflow pipeline that converts gVCF files into Log R Ratio and B Allele Frequency signals compatible with established CNV callers. Applied to 12,509 individuals from the SPARK cohort, gVCF2CNV generated signals at an average of 2.7 million SNV positions per individual and completed signal extraction in 4 hours using 192 CPUs. CNV calling with PennCNV and QuantiSNP identified candidate CNVs across a broad size range, with trio-based Mendelian precision reaching approximately 80% or higher for deletions of at least 30 kb and duplications of at least 5 kb. Application to 414,824 individuals from the All of Us cohort was completed in 96 hours, demonstrating feasibility at biobank scale. These results show that gVCF files can serve as a scalable input for CNV detection in large WGS cohorts.

**Availability and Implementation:** gVCF2CNV is available at https://github.com/JacquemontLab/gVCF2CNV, implemented as a Nextflow pipeline with Perl and Python components, supported on Linux.

## INTRODUCTION

Copy-number variants (CNVs) are an important class of structural variation and contribute to human disease, neurodevelopmental disorders, and quantitative trait variation (Salgado *et al*. 2020). Large CNVs are routinely detected using genotyping arrays (Wang *et al*. 2007) and have been widely studied in population and disease cohorts. In contrast, smaller CNVs, including intragenic events, remain more difficult to characterize at scale because their detection requires higher-resolution genomic data.

Whole-genome sequencing (WGS) provides higher genomic resolution than genotyping arrays and is now available in large cohorts (Bick *et al*. 2024). However, CNV detection from WGS remains computationally expensive at scale. Most WGS-based CNV callers require BAM or CRAM files as input, which are costly to store, often difficult to access in controlled-access cohorts, and computationally intensive to process. A recent large-scale WGS CNV analysis of 470,727 UK Biobank genomes required 72 days and 7.3 million CPU hours for BAM-based calling and focused on CNVs larger than 10 kb (Zou *et al*. 2026).

Genomic VCF (gVCF) files represent a practical alternative. They are routinely produced by standard variant-calling workflows and contain genotype, read depth, allele depth, and genotype-quality information. Compared with BAM or CRAM files, gVCFs are smaller, easier to distribute within controlled-access environments, and already available in many WGS studies. However, gVCF files are not directly compatible with widely used CNV callers such as PennCNV (Wang *et al*. 2007a) and QuantiSNP (Colella *et al*. 2007) which were originally developed to use Log R Ratio (LRR) and B Allele Frequency (BAF) signals derived from SNP-array intensities. LRR reflects relative copy number signal, whereas BAF reflects allelic balance. The read depth and allele-depth information available in gVCF files provides an opportunity to derive analogous LRR and BAF signals from sequencing data.

Here, we present gVCF2CNV, a modular Nextflow pipeline that converts individual gVCF files into sequencing-derived LRR and BAF signals compatible with established CNV callers. We evaluate gVCF2CNV in 12,509 individuals from the SPARK cohort, including 1,103 parent-offspring trios, and demonstrate its scalability by applying it to 414,824 individuals from the All of Us cohort. Together, these analyses show that gVCF2CNV enables large-scale CNV detection from gVCF files when BAM/CRAM-based processing is impractical.

## MATERIALS AND METHODS

### Dataset

We analyzed 30× whole-genome sequencing data from 12,509 individuals in the SPARK cohort (SPARK Consortium 2018), including 3,314 individuals with autism spectrum disorder and 1,103 parent-offspring trios. The dataset included individual gVCF files generated using DeepVariant, chromosome-level population VCF files, and phenotypic information. Sequencing data were aligned to the GRCh38 reference genome.

### gVCF2CNV pipeline overview

gVCF2CNV is implemented as a modular Nextflow DSL2 pipeline executed on a SLURM high-performance computing cluster (Figure 1). The pipeline consists of three main steps: (i) construction of a cohort-level SNV reference panel from population VCF files, (ii) extraction of LRR and BAF signals from individual gVCF files, and (iii) construction of a cohort-specific PFB file. The pipeline generates individual LRR/BAF files and a cohort-specific PFB file, which serve as input for established LRR/BAF-based CNV callers such as PennCNV and QuantiSNP.

**Figure 1.**
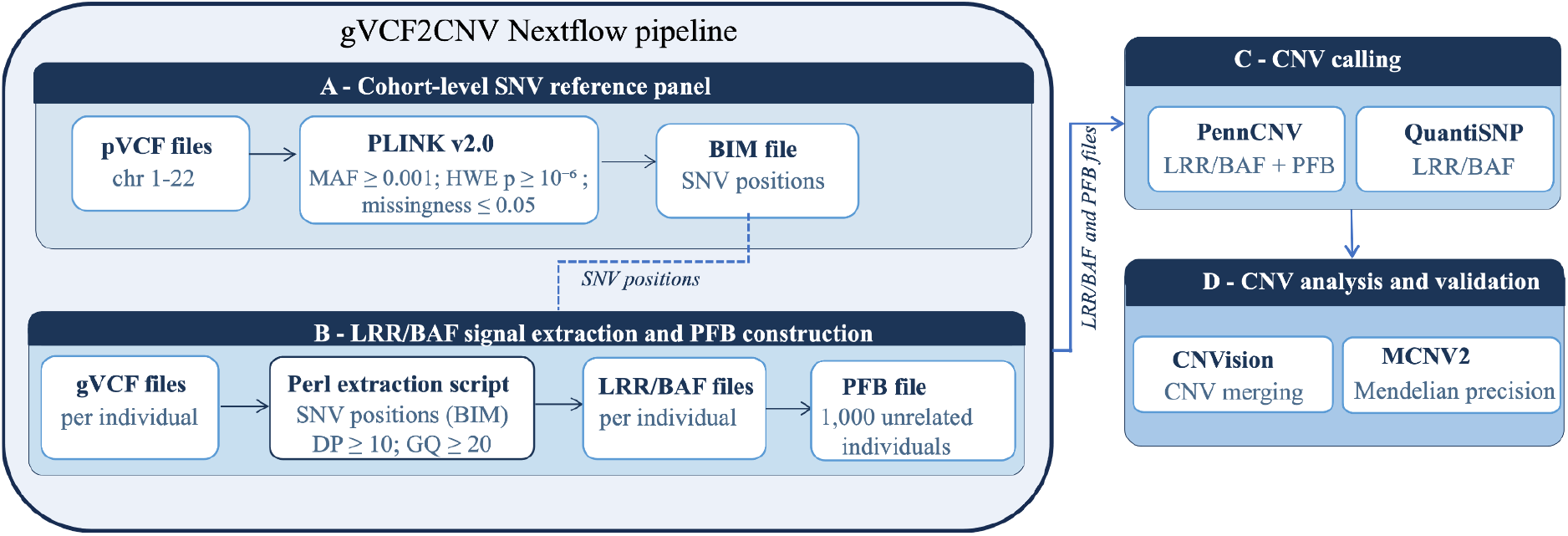
Overview of the gVCF2CNV pipeline and downstream CNV analysis. (A) Cohort-level SNV reference panel construction. Population VCF files for chromosomes 1-22 are converted to PLINK format and filtered based on minor allele frequency (MAF ≥0.001), Hardy–Weinberg equilibrium (p ≥1 × 10-6), missingness (≤0.05), and overlap with problematic genomic regions. The retained SNV positions are stored in a BIM file, which serves as the cohort-level reference panel for subsequent LRR/BAF extraction. (B) LRR/BAF extraction and PFB construction. For each individual, LRR and BAF are generated from gVCF records at retained SNV positions meeting read depth (DP ≥10) and genotype quality (GQ ≥20) thresholds using a Perl extraction script. A cohort-specific population frequency of B allele (PFB) file is constructed using 1,000 unrelated individuals. Steps A and B constitute the gVCF2CNV Nextflow pipeline. (C) CNV calling. LRR/BAF files and the cohort-specific PFB file are used as input for PennCNV, whereas QuantiSNP uses LRR/BAF files only. (D) CNV analysis and validation. Calls from both algorithms are compared and merged using CNVision, including assessment of reciprocal overlap, and subsequently evaluated using the MCNV2 framework for Mendelian precision analysis.

### Construction of the cohort-level SNV reference panel

For each of the 22 autosomes, population VCF files were converted to PLINK format using PLINK v2.0 and subsequently merged. Standard genotype quality-control filtering included variant and individual missingness, Hardy-Weinberg equilibrium, and minor allele frequency thresholds (Marees *et al*. 2018). Samples with genotype missingness >5% were excluded. Biallelic SNVs were retained if variant missingness was ≤5%, Hardy-Weinberg equilibrium p ≥1×10-6, and minor allele frequency was ≥0.001. Variants overlapping problematic genomic regions were excluded. These regions were defined using a curated annotation file combining segmental duplications, centromeres, telomeres, the major histocompatibility complex, and problematic-region tracks from UCSC (Kent *et al*. 2002), retrieved in April 2025. Overlapping intervals were merged using BEDTools. The annotation file, the source URLs, and the code used to generate it are distributed with the pipeline for both GRCh37 and GRCh38. After filtering, the final PLINK BIM file contained approximately 13.6 million SNV positions and was used as the cohort-level SNV reference panel.

### Extraction of LRR and BAF signals from gVCF files

LRR and BAF signals were extracted from individual gVCF files using a Perl script implemented in gVCF2CNV and adapted from convertMapToSignal.pl, originally developed for PennCNV-SEQ (Araújo Lima de and Wang 2017). The original script extracts read depth and allele counts from BAM files using SAMtools and BCFtools (Danecek *et al*. 2021). In gVCF2CNV, these values are obtained directly from the read depth (DP) and allele-depth (AD) fields of gVCF records, removing the need to process alignment files. Non-variant reference blocks were not used for signal generation.

For each individual, signal extraction was restricted to biallelic SNV positions included in the cohort-level SNV reference panel. Sites were retained if they met user-defined thresholds for read depth (DP) and genotype quality (GQ), set here to DP ≥10 and GQ ≥20. GQ reflects the confidence in the genotype call. Homozygous-reference calls were excluded. For each retained SNV, BAF and LRR were computed as:

BAF = AD_alt / DP

LRR = log(DP_i / mean DP)

where AD_alt is the allele depth of the alternative allele, DP_i is the total read depth at position i, and mean DP is the mean read depth across all retained SNVs for that individual. After filtering, an average of 2.7 million SNV positions per individual were retained for signal generation.

### Construction of the PFB file

A cohort-specific PFB file was constructed using BAF data from 1,000 unrelated individuals. Relatedness was estimated using KING (Manichaikul *et al*. 2010), and individuals with lower genotype missingness were prioritized for inclusion. For each SNV, the PFB value was calculated as the mean BAF across selected individuals for whom the position passed quality filters. PFB computation was implemented in Python using the Polars library. The final PFB file contained 13,070,685 SNVs.

### CNV calling and quality control

Individual LRR/BAF files were used as input for PennCNV (Wang et al. 2007) and QuantiSNP (Colella et al. 2007), two HMM-based CNV callers originally developed for SNP-array data. The cohort-specific PFB file was additionally provided to PennCNV. For PennCNV, we used the HMM file provided by PennCNV-SEQ (Araújo Lima de and Wang 2017). CNV calls were generated for all 12,509 SPARK individuals. Calls from both algorithms were merged using CNVision (Sanders *et al*. 2011). A minimum confidence score of 30 was applied to retained CNVs. CNVs overlapping problematic genomic regions by more than 50% were excluded from downstream analyses.

Sample quality was assessed using three metrics reported by PennCNV: LRR standard deviation (LRR_SD), waviness factor (WF), and BAF drift, which capture genome-wide variability in read depth, systematic depth fluctuations along chromosomes, and deviations in BAF distributions, respectively. Samples were excluded if they exceeded any of the following thresholds recommended by Wang et al. (2007): LRR_SD > 0.35, |WF| > 0.05, or BAF drift > 0.01.

### CNV annotation

CNVs were annotated with gene content using the annotation module of MCNV2 (Diop *et al*. 2026), based on Ensembl gene coordinates for GRCh38.

### CNV validation: Mendelian precision

CNV call quality was evaluated in 1,103 parent-offspring trios using Mendelian precision (MP) (Diop *et al*. 2026). MP is defined as the proportion of offspring CNVs also detected in at least one parent. Before MP computation, CNVs overlapping problematic genomic regions by more than 50% were excluded. CNVs encompassing at least one haploinsufficient gene (LOEUF <0.6) were also excluded to reduce the contribution of potentially true *de novo* events to the MP estimate. An offspring CNV was classified as inherited when a parental CNV of the same type showed at least 50% reciprocal overlap with the offspring CNV. MP was computed across nine size categories: 1-5 kb, 5-10 kb, 10-30 kb, 30-50 kb, 50-100 kb, 100-200 kb, 200-500 kb, 500 kb-1 Mb and ≥1 Mb, and stratified by CNV type, caller, and confidence score threshold.

### Scalability: All of Us cohort

We applied gVCF2CNV to 414,824 individuals from the All of Us cohort to evaluate scalability at the biobank scale. LRR and BAF signals were generated using the same approach as for SPARK, yielding an average of 3.5 million positions per sample. CNV calling was performed using PennCNV only. Standard quality filters were applied, including a minimum confidence score of 30, a minimum CNV length of 1 kb, and exclusion of CNVs overlapping more than 50% with problematic genomic regions.

### Implementation

gVCF2CNV is available at https://github.com/JacquemontLab/gVCF2CNV. Signal extraction is implemented in Perl and PFB construction in Python using Polars. The pipeline runs on SLURM clusters using Nextflow DSL2. Software versions are documented in the repository.

## RESULTS

### gVCF2CNV generated LRR/BAF signals from SPARK gVCF files

We applied gVCF2CNV to 12,509 individuals from the SPARK cohort using the workflow shown in Figure 1. After population-level VCF quality control and exclusion of problematic genomic regions, the cohort-level SNV reference panel contained approximately 13.6 million SNVs (Figure 1A). At the individual level, a mean of 2.7 million SNVs passed filtering and were retained for LRR/BAF generation (Figure 1B). The final PFB file contained 13,070,685 SNVs. Individual LRR/BAF files were used for CNV calling with PennCNV and QuantiSNP, with the cohort-specific PFB file additionally provided to PennCNV (Figure 1C).

LRR/BAF extraction for all 12,509 individuals was completed in 4 hours using 192 CPUs. This corresponds to approximately 768 CPU-hours in total, or approximately 0.061 CPU-hours per individual.

These results show that gVCF2CNV can generate LRR and BAF signals from gVCF files at cohort scale, without requiring access to BAM or CRAM files.

### WGS-derived LRR/BAF signals showed stable sample-level quality metrics

We assessed sample-level signal quality using LRR standard deviation, waviness factor, and BAF drift (Figure 2A-C). These three metrics assess genome-wide variability in read depth, systematic depth fluctuations along chromosomes, and the stability of BAF distributions.

**Figure 2.**
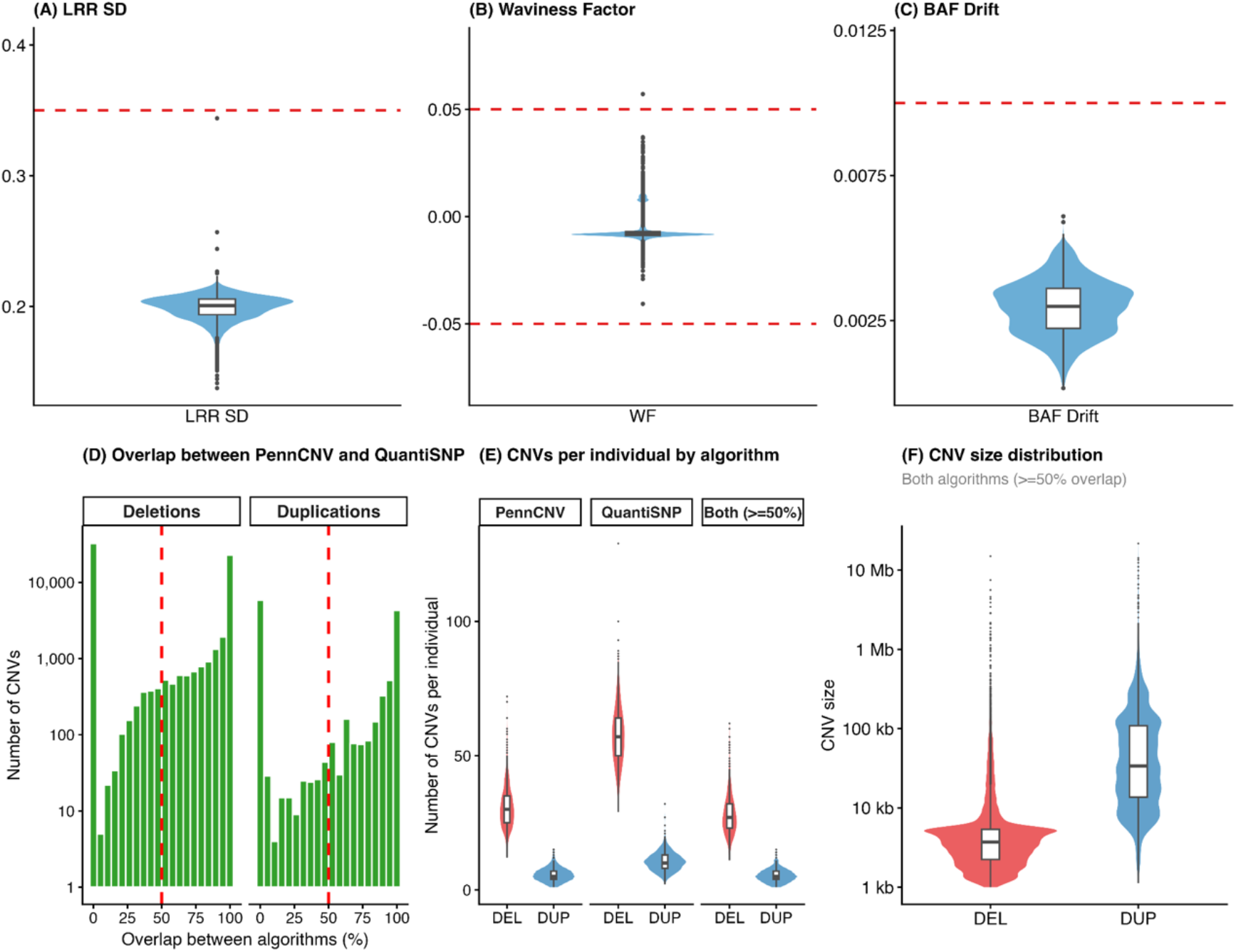
Quality control metrics and CNV distribution. (A-C) Signal quality control across all 12,509 individuals. Distributions of LRR standard deviation (A), waviness factor (B), and BAF drift (C). Red dashed lines indicate the quality-control thresholds used in this study (LRR_SD ≤0.35, |WF| ≤0.05, BAF drift ≤0.01). (D-F) CNV distributions in offspring from 1,103 complete parent-offspring trios, after applying quality filtering, including a confidence score ≥30, exclusion of CNVs overlapping problematic genomic regions by more than 50%, and exclusion of the sample failing the WF threshold. (D) Distribution of reciprocal overlap between PennCNV and QuantiSNP calls. The red dashed line indicates the 50% reciprocal-overlap threshold used to define concordant calls. (E) Number of CNVs per individual detected by PennCNV, QuantiSNP, or both algorithms (≥50% reciprocal overlap), stratified by deletion (DEL, red) and duplication (DUP, blue). (F) Size distribution of deletions and duplications identified by both algorithms (≥50% reciprocal overlap).

Using thresholds of LRR standard deviation ≤0.35, absolute waviness factor ≤0.05, and BAF drift ≤0.01, no samples failed the LRR standard deviation threshold (Figure 2A), one sample exceeded the waviness-factor threshold (Figure 2B), and all samples passed the BAF drift threshold (Figure 2C). These results indicate that gVCF-derived LRR/BAF signals were stable across most individuals in the SPARK cohort (Figure 2).

### CNV calling identified candidate CNVs across a broad size range

CNV calling was performed using PennCNV and QuantiSNP, followed by merging with CNVision (Figure 1C, D). Across 12,509 individuals, 1,388,045 unfiltered CNVs were detected. These included 1,167,465 deletions and 220,580 duplications.

We applied the following quality filters: exclusion of one sample failing the WF threshold, confidence score ≥30, and exclusion of CNVs overlapping problematic genomic regions by more than 50%. Across the full SPARK cohort, 812,827 candidate CNVs were retained: 715,814 deletions (88%) and 97,013 duplications (12%). Most candidate CNVs were smaller than 50 kb, with the largest number of calls in the 1-5 kb size category.

The size distribution differed between deletions and duplications. Deletions were more common among CNVs below 50 kb, whereas duplications represented a larger proportion of calls above 50 kb.

### PennCNV and QuantiSNP showed partial caller concordance

We next evaluated the overlap between CNVs detected by PennCNV and QuantiSNP in 1,103 offspring from complete parent-offspring trios. Among this subset, 76,699 offspring CNVs were retained, including 64,844 deletions and 11,855 duplications.

PennCNV detected 40,244 CNVs and QuantiSNP detected 74,862 CNVs. A total of 36,501 (48%) CNVs were concordant between algorithms, with at least 50% reciprocal overlap. Among concordant CNVs, individuals carried a median of 27 deletions (IQR 22-32) and 3 duplications (IQR 2-5) (Figure 2E). Deletions showed a median size of 3.7 kb (IQR 2.2-5.4 kb) and duplications a median of 34 kb (IQR 14-109 kb) (Figure 2F).

The reciprocal-overlap distribution was bimodal, with many events showing either minimal or near-complete overlap between callers (Figure 2D). This caller comparison suggests that PennCNV and QuantiSNP identify a shared set of high-confidence events while also producing caller-specific calls, particularly among smaller CNVs. We therefore report caller-specific and concordant-call results separately where possible.

### Trio-based Mendelian precision increased with CNV size and confidence score

We used parent-offspring trios to assess inheritance consistency of candidate CNV calls. Mendelian precision (MP) was calculated as the proportion of offspring CNVs observed in at least one parent. Because true *de novo* CNVs occur in parent-offspring trios, especially among larger and clinically relevant CNVs, Mendelian precision should be interpreted as an inheritance-consistency metric rather than a direct estimate of false-discovery rate.

CNVs encompassing at least one haploinsufficient gene (LOEUF <0.6) were excluded before MP computation, to separate detection accuracy from biological *de novo* rate. MP was computed across nine size categories, stratified by confidence score threshold, and separately for CNVs detected by PennCNV, QuantiSNP, and both algorithms with at least 50% reciprocal overlap (Figure 3).

**Figure 3.**
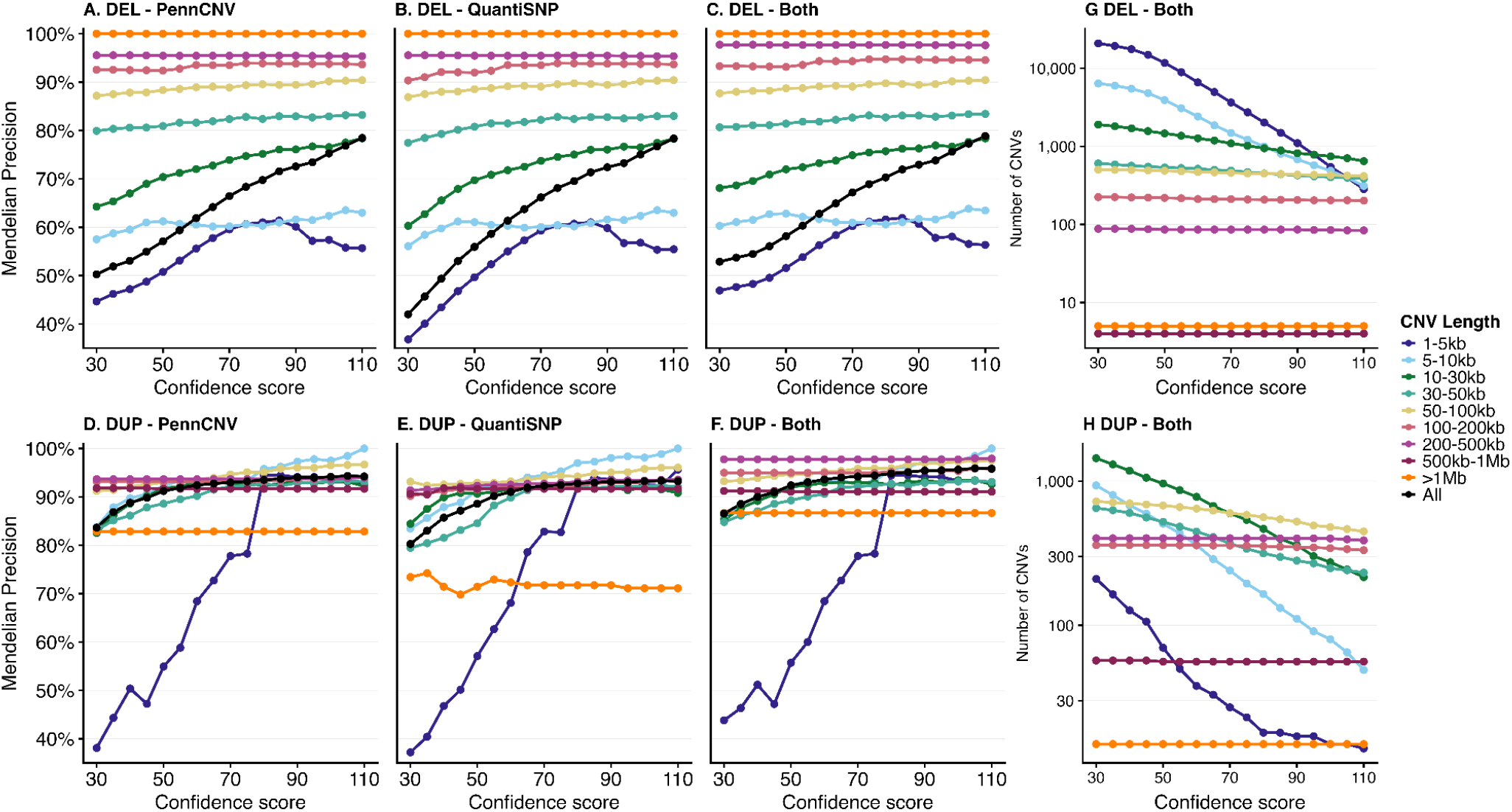
Mendelian Precision (MP, %) of CNVs as a function of minimum confidence score threshold across size categories. At each threshold value shown on the x-axis, only CNVs with a confidence score equal to or above that threshold are included in the MP computation (e.g., x = 50 means CNVs with score ≥50). MP for deletions (top row) and duplications (bottom row) detected by PennCNV (A, D), QuantiSNP (B, E), and CNVs detected by both algorithms with at least 50% reciprocal overlap (C, F). Panels G and H show the number of CNVs detected by both algorithms retained at each confidence score threshold for deletions and duplications, respectively. Each color-coded line corresponds to a size category: 1-5 kb, 5-10 kb, 10-30 kb, 30-50 kb, 50-100 kb, 100-200 kb, 200-500 kb, 500 kb-1 Mb and ≥1 Mb. The black curve represents MP across all size categories combined. Prior to MP computation, CNVs overlapping problematic genomic regions by more than 50% were excluded, as were CNVs encompassing at least one haploinsufficient gene (LOEUF < 0.6).

For deletions, MP increased with both CNV size and confidence-score threshold (Figure 3A, B, C). Deletions in the 1-5 kb range showed the lowest Mendelian precision, rising from approximately 45% at a score of 30 to a peak near 62% before declining at the highest thresholds. Deletions in the 5-10 kb and 10-30 kb ranges reached approximately 63% and 78%, respectively. Deletions of 30-50 kb approached or exceeded 80%, and deletions above 50 kb generally exceeded 80% across confidence thresholds, and deletions larger than 1 Mb showed near-complete inheritance consistency. The number of deletions retained at each confidence score decreased sharply for the smallest size category, from over 20,000 at a score of 30 to a few hundred at a score of 110, while larger size categories remained comparatively stable (Figure 3G).

For duplications, MP was approximately 80% or higher for most size categories above 5 kb, even at the lowest confidence threshold (Figure 3D, E, F). Duplications in the 1-5 kb range showed the lowest values, rising from approximately 38% at a score of 30 to over 90% at higher thresholds, but this came at the cost of retaining very few CNVs (Figure 3H). Duplications larger than 1 Mb showed lower inheritance consistency than intermediate-size categories, potentially reflecting a higher contribution of true *de novo* events.

Restricting to CNVs detected by both PennCNV and QuantiSNP produced Mendelian precision estimates comparable to single-caller results for several size categories (Figure 3C, F).

Together, these results support the use of gVCF2CNV for cohort-scale CNV detection, particularly when analyses are stratified by CNV size, type, confidence score, and caller concordance.

### gVCF2CNV scaled to 414,824 All of Us genomes

To evaluate scalability beyond SPARK, we applied gVCF2CNV to 414,824 individuals from the All of Us cohort. CNV calling was performed using PennCNV only. The analysis was completed in 96 hours of wall-clock time using approximately 450,000 CPU hours.

Compared with BAM-based CNV calling at similar cohort scale, the gVCF-based workflow substantially reduced the computational burden and avoided direct processing of BAM or CRAM files. However, this comparison should be interpreted as a practical scalability benchmark rather than a matched accuracy benchmark, because the methods differ in input data, signal type, caller design, and CNV size range.

These results demonstrate that gVCF2CNV can be applied to biobank-scale WGS datasets in controlled-access environments where gVCF files are available, but BAM/CRAM processing is computationally or logistically prohibitive.

## DISCUSSION

We developed gVCF2CNV to address a practical gap in large-scale WGS CNV analysis: many cohorts provide gVCF files, whereas BAM/CRAM-based CNV calling remains computationally expensive and logistically difficult. By converting gVCF records into LRR and BAF signals, the pipeline allows established LRR/BAF-based CNV callers to be applied to WGS-derived data. In SPARK, gVCF2CNV generated LRR/BAF signals at 2.7 million SNV positions per individual in 4 hours using 192 CPUs. CNV calling with PennCNV and QuantiSNP produced candidate CNV calls whose trio-based Mendelian precision increased with size and confidence score. Application to All of Us further demonstrated that the approach can scale to hundreds of thousands of genomes.

The central motivation for gVCF2CNV is that gVCFs are already generated by many WGS variant-calling pipelines and contain information that is informative for CNV detection, including genotype, read depth, allele depth, and genotype quality (Poplin *et al*. 2018). In contrast to BAM or CRAM files, gVCFs are smaller and more portable within controlled-access environments. gVCF2CNV therefore shifts CNV screening from alignment-level reprocessing to variant-record-level signal extraction. This design is particularly useful for large biobank and disease cohorts in which access to alignment files is restricted, storage costs are high, or reprocessing raw alignments is impractical.

gVCF2CNV is complementary to existing CNV callers. Algorithms such as PennCNV and QuantiSNP use LRR and BAF signals and have been widely applied to SNP-array data. WGS-based CNV callers can use richer evidence, including read depth, read-pair orientation, split reads, and local assembly, and can provide more precise breakpoint resolution (Kosugi *et al*. 2019). gVCF2CNV occupies an intermediate space: it does not attempt to infer CNVs directly from raw alignments, but instead generates WGS-derived LRR/BAF signals that can be processed by established LRR/BAF-based callers. This makes the approach attractive for scalable screening, while retaining the need for orthogonal validation in applications requiring exact breakpoints or clinical interpretation.

The high resolution of WGS-derived SNV positions provides substantially more potential signal locations than standard genotyping arrays, supporting candidate CNV detection below the size range typically targeted by array-based CNV pipelines. In SPARK, most candidate events were smaller than 50 kb, including many events in the 1-5 kb range. These small calls should be interpreted with caution. Trio-based Mendelian precision improved with increasing confidence score and CNV size, indicating that the smallest events are more sensitive to thresholding and caller behavior. For downstream analyses, size-stratified filtering and reporting are therefore essential, particularly when studying intragenic or noncoding CNVs.

Trio-based Mendelian precision provides a useful validation framework because inherited CNVs should be observed in an offspring and at least one parent (Diop *et al*. 2026). In this study, Mendelian precision increased with CNV size and confidence score, supporting the utility of the generated LRR/BAF signals. However, Mendelian precision is not equivalent to truth-set precision. True *de novo* CNVs reduce apparent inheritance consistency, which may contribute to the lower values observed for the largest duplications (Sanders *et al*. 2011). Requiring 50% reciprocal overlap may underestimate concordance when parent and offspring calls capture the same event with slightly different boundaries. Future benchmarking against orthogonal WGS CNV callsets, matched SNP-array data, curated recurrent CNVs, or simulated events will be needed to quantify sensitivity and false-discovery rates directly (Zook *et al*. 2020).

Several limitations define the current scope of gVCF2CNV. First, the current implementation uses retained variant positions and does not yet exploit non-variant reference blocks, which may reduce signal continuity in regions with low SNV density. Second, the depth filter improves signal stability but limits detection of homozygous deletions, because very-low-depth sites are excluded before signal generation. Third, PennCNV and QuantiSNP (Colella *et al*. 2007, Wang *et al*. 2007) were originally parameterized for SNP-array signals, and further calibration may improve performance on WGS-derived LRR/BAF data. Finally, the current study emphasizes scalability and trio-based inheritance consistency; full accuracy benchmarking against orthogonal callsets remains an important next step.

In summary, gVCF2CNV provides a practical route for CNV screening in large WGS cohorts where gVCF files are available, but BAM/CRAM processing is infeasible. By converting gVCF records into LRR and BAF signals compatible with established CNV callers, the pipeline lowers the computational barrier to copy number variation analysis and enables size-stratified CNV discovery at cohort scale.

## Funding

This work was supported by a grant from the National Institutes of Health: 5U01MH119690 (SJ) and the Natural Sciences and Engineering Research Council of Canada through an Individual Discovery Grant RGPIN-2025-04416 and a Université de Montréal Department Chair Grant (SH).

## Acknowledgements

This research was enabled by support provided by the Digital Research Alliance of Canada (https://www.alliancecan.ca/). We are grateful to all of the families in SPARK, the SPARK clinical sites, and SPARK staff. We appreciate obtaining access to phenotypic and genetic data on SFARI Base. Approved researchers can obtain the SPARK population dataset described in this study by applying at https://base.sfari.org.

## Notes

### Competing Interest Statement

The authors have declared no competing interest.

